# Replication protein Rep provides selective advantage to viruses in the presence of CRISPR-Cas immunity

**DOI:** 10.1101/2021.11.18.469202

**Authors:** Weijia Zhang, Yuvaraj Bhoobalan-Chitty, Xichuan Zhai, Yan Hui, Lars Hestbjerg Hansen, Ling Deng, Xu Peng

## Abstract

Prokaryotic viruses express anti-CRISPR (Acr) proteins to inhibit the host adaptive immune system, CRISPR-Cas. While the virus infection biology was shown to be strongly dependent on the relative strengths of the host CRISPR-Cas and viral Acrs, little is known about the role of the core processes of viral life cycle (replication, packaging etc) in defence/anti-defence arms race. Here, we demonstrate the selective advantage provided by a replication initiator, Rep, in the context of CRISPR-Acr interactions. First, we developed a two-host based CRISPR-Cas genome editing tool for the deletion of highly conserved and thus potentially important viral genes. Using this strategy, we deleted a highly conserved Rep-coding gene, *gp16*, from the genome of *Sulfolobus islandicus* rod-shaped virus 2 (SIRV2). The knockout mutant (Δ*gp16*) produced around 4 fold less virus in a CRISPR-null host, suggesting that Rep is the major contributor to replication initiation in Rudiviridae. Indeed, DNA sequencing revealed Rep-dependent replication initiation from the viral genome termini, in addition to Rep-independent replication initiation from non-terminal sites. Intriguingly, the lack of Rep showed a profound effect on virus propagation in a host carrying CRISPR-Cas immunity. Accordingly, the co-infecting parental virus (*rep*-containing) outcompeted the Δ*gp16* mutant much more quickly in CRISPR-containing host than in CRISPR-null host, demonstrating a selective advantage provided by Rep in the presence of host CRISPR-Cas immunity. Despite the non-essentiality, *rep* is carried by all known members of Rudiviridae, which is likely an evolutionary outcome driven by the ubiquitous presence of CRISPR-Cas in Sulfolobales.

**Importance:** CRISPR-Cas and anti-CRISPR proteins are accessary to prokaryotes and their viruses respectively. To date, research has been focused on their diversity, molecular mechanisms and application in genome editing. How CRISPR-Acr arms race influence the evolution of viral core genes involved in the basic virus life cycle remained a gap of knowledge so far. This study provides the first evidence that CRISPR-Acr arms race poses a selection pressure on the efficiency of viral genome replication, forcing viruses to evolve highly productive replication machineries..

## INTRODUCTION

All life forms and their viruses are subject to continuous evolution, driven by the need to gain an advantage over the other in order to avoid complete extinction. Among the diverse antiviral defense systems, clustered regularly interspaced small palindromic repeats (CRISPR) and CRISPR-associated (Cas) genes constitute the only adaptive immune system in prokaryotes which is present in 40% bacteria and 90% archaea, and classified into two classes, six types and 33 subtypes [1]. To counteract the CRISPR-Cas immunity, bacteriophages and archaeal viruses have evolved a diverse range of anti-CRISPR (Acr) proteins that inhibit CRISPR-Cas at different stages [2, 3]. To date, 88 Acr families have been identified [4, 5] which exhibit diverse inhibition mechanisms and structures [2].

While the high diversity of CRISPR-Cas and Acrs has been appreciated and their molecular mechanisms have been extensively studied, insights into the impact of CRISPR-Acr interactions on bacteriophage infection biology are just emerging. On one hand, Acr phages cooperate such that accumulating Acr proteins expressed from targeted phage genome inhibit the host CRISPR-Cas immunity, enabling subsequent infection to complete in the immunosuppressed host. This cooperation requires the viral population to be above a threshold multiplicity of infection (MOI) which was found to be inversely proportional to the Acr “strength” (i.e., the efficiency of immunosuppression). On the other hand, the strength of host CRISPR-Cas immunity directly impacts the MOI threshold, e.g. a higher MOI threshold is related to a higher number of targeting CRISPR spacers [6, 7].

Despite recent advances in elucidating CRISPR-Acr interactions, very little is known about the possible interplay between viral core life cycle processes (e.g. replication, gene expression and assembly) and the defense/anti-defense arms race. In this work, we studied the significance of a replication protein (Rep) encoded by *Sulfolobus islandicus* rod-shaped virus 2 (SIRV2) in relation to CRISPR-Acr combat. SIRV2 is a lytic archaeal virus whose entry [8, 9], transcription regulation [10, 11], replication [12–15], virion release [16] and Acrs [17–20] have been studied. Moreover, gene editing tools are available for deleting, inserting, or mutating SIRV2 genes [21, 22]. The virus carries a linear dsDNA genome of 35 kbp with inverted terminal repeats (ITR) of 1.6 kbp and the two strands are covalently linked at the termini [23]. The hairpin terminal structure, together with biochemical and microscopic data, led to the following replication model: nicks are made at the genomic termini allowing multiple replication initiation from a single genome template, followed by strand displacement replication; the multiple ssDNA replicative intermediates remain attached to the genome template until 4 hours post infection (hpi) forming a brush-like structure [12–14, 23]. Furthermore, in addition to strand displacement replication, bidirectional strand-coupled replication was inferred to take part in SIRV2 replication. However, the latter only contributes to the generation of a minor fraction of the replicative intermediates [13]. The lytic life cycle spans about 12 – 13 hours [16].

SIRV2 *gp16* encodes a protein belonging to the replication initiator (Rep) superfamily which contains protein members responsible for initiation of rolling circle replication in diverse viruses and plasmids [24, 25]. The SIRV Rep was shown *in vitro* to nick a ssDNA oligo carrying the sequence of the viral genomic termini and form a covalent adduct with the 5’ end of the cleaved strand [12]. This provided evidence for the SIRV Rep being a replication initiator. In this study, we provide *in vivo* evidence that SIRV2 Rep is responsible for replication initiation at the genomic termini. More importantly, Rep provides replicative advantage in the presence of host CRISPR-Cas immunity.

## Material and Methods

### Strains and growth conditions

*Sulfolobus islandicus* LAL14/1 and its derivatives were grown aerobically at 78°C and 150 rpm in SCV medium (basic salts medium supplemented with 0.2% sucrose, 0.2% casamino acids, and a vitamin mixture [26]) or SCVU (SCV supplemented with uracil of 20 μg/mL, final concentration). *E*.*coli* DH5α was used for cloning of the *Sulfoloubs*-*Escherichia* shuttle plasmids [27].

### Plasmid construction, PCR and Primers

PCRs were performed with Phusion Hot Start II DNA Polymerase (Thermo Fisher Scientific) or Phanta Max Super-Fidelity DNA Polymerase (Vazyme) using primers listed in Table S1. Plasmids carrying CRISPR spacers and homologous sequences for recombination were constructed using protocols described earlier [28, 29]. For construction of the complementation plasmid encoding a mutated *gp16*, a silent mutation was introduced by overlap extension PCR. Additional information about primers, restriction sites and vectors used are listed in Table S1.

### Genetic manipulation of viruses

Deletion of *gp16* was performed using a CRISPR-based genome-editing method described earlier [21, 22], except that two engineered hosts were used sequentially in this case (Figure 1A). Briefly, plasmids for the engineering of the two hosts were constructed using primers listed in Table S1, purified and transformed by electroporation [22] into *S. islandicus* LAL14/1 Δarrays [29].

**Figure 1.**
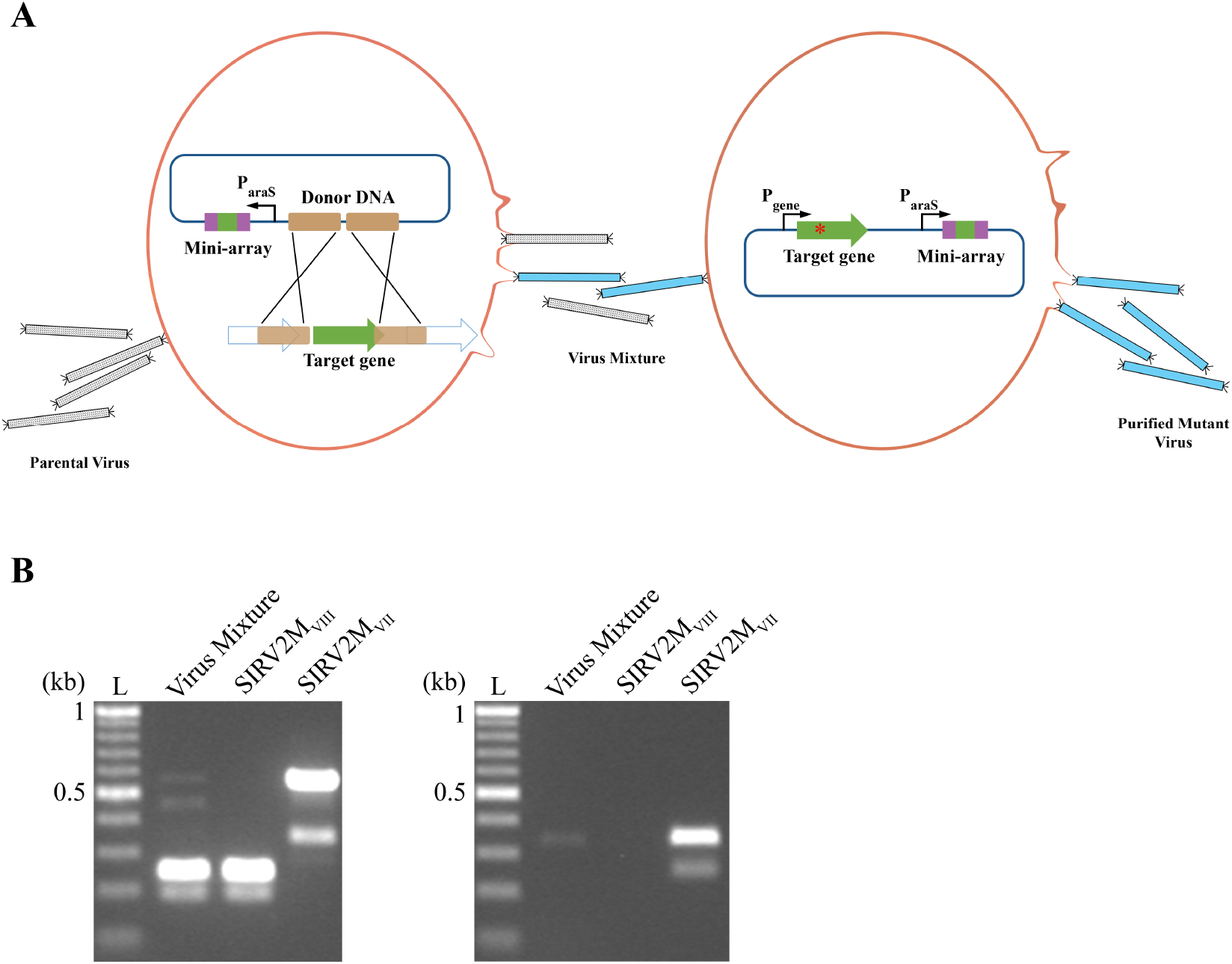
Deletion of the highly conserved *gp16*. (A) Schematic illustration of the method. The deletion strain carrying the genome editing plasmid was infected with the parental virus (grey rods). Part of the viral genome in the deletion strain is depicted showing the target gene *gp16* (green arrow) and the flanking genes (empty arrows). Recombination arms on the viral genome and the genome editing plasmid are indicated with brown blocks. A CRISPR mini-array containing two repeats (purple blocks) and a spacer (green block) is under the control of an arabinose promoter (black arrow). Infection of the deletion strain with the parental virus generates a mixture of *Δgp16* (blue rods) and the parental virus. The purification strain carries a plasmid encoding the same CRISPR mini-array as that in the deletion strain and a *gp16* gene with silent mutations (red dot). (B) PCR analysis of the infected culture supernatant demonstrating the deletion of *gp16* from SIRV2M_VII_. Origins of PCR templates are shown on top. Left panel, primers base pairing with *gp16* flanking sequences were used in the PCR. L: DNA ladder with sizes (kb) shown at left. Virus mixture: supernatant from the infected deletion strain; SIRV2M_VIII_: supernatant from the infected purification strain; SIRV2M_VII_: parental virus particles. Right panel, same as the left panel except that primers specific for the deleted region were used in PCR. Samples were identical to those in the left panel. Lane 3 in both panels represents the parental virus (SIRV2M_VII_). Primers used in these experiments are listed in Table S1.

### Cell growth and virus titer of infected cultures

*S. islandicus* LAL14/1 Δarrays carrying either an empty vector (Δarrays/pcontrol) or a plasmid encoding SIRV2 gp16 (Δarrays/p*gp16*) were grown to an OD_600_ between 0.05-0.1 and infected with a virus (Δ*gp16*-a or parental virus) at the specified MOIs. The wild-type *S. islandicus* LAL14/1 carrying a complementation plasmid (p*acrIIIB2*) was infected at OD_600_ 0.05 with Δ*gp16*-b or its corresponding parental virus at an MOI of 10^−5^. For competition experiments, cells were infected with a virus mixture of Δ*gp16*-b and its corresponding parental virus at a ratio of 10:1. For infection of the strain carrying only CRISPR-Cas subtype III-B system, the MOI of Δ*gp16*-b was set at 0.025 and accordingly the MOI of the parental virus was 0.0025; for infection of the remaining strains (Figure 5), an MOI of 10^−5^ was used for Δ*gp16*-b and 10^−6^ for the parental virus. Cell growth was monitored by measuring OD_600_ at specific intervals over several days as indicated in the respective figures. Infected cultures were sampled at appropriate time points for virus titration by plaque assay and for quantification of virus strains by PCR. Three biological replicates were performed in all experiments.

### Quantification of intracellular viral DNA using quantitative PCR (qPCR)

Prior to infection, the Δ*gp16*-a mutant was purified and concentrated by polyethylene glycol precipitation to obtain a high virus titer [13]. Δarrays/pcontrol and Δarrays/p*gp16* cells of 200 mL were grown to an OD_600_ of 0.2 and infected with Δ*gp16*-a at an MOI of 0.1. Samples of 15 mL were taken at 0, 0.5, 1, 2, 4, 6 and 8 hpi, centrifuged at 6,300 x g. Virus titer of the supernatants was determined by plaque assay (see below). Cell pellets were resuspended in 194 μL basic medium salt solution to which DNase I (#EN0521, Thermo Fisher Scientific) was added at 0.05 units/μL along with 5 μL of 10x reaction buffer, incubated at 37°C for 1 h to remove contaminating extracellular DNA and the DNase was inactivated afterwards at 65°C for 10 min. DNA was then extracted using the DNeasy Blood & Tissue Kit (QIAGEN N.V., Hilden, Germany) as per the manufacturer’s instructions. Viral DNA was extracted from SIRV2 virion by phenol-chloroform method followed by ethanol precipitation. All DNA samples were diluted to the same DNA concentration (40 ng/μL), and serial dilutions of virion DNA were used for constructing the standard curve. Primer pairs specific to *gp01, gp26* and *gp38* (Table S1) were designed to quantify replication at different locations on the viral genome. qPCR reactions were performed in 10 μL mixtures, containing 5 μL PowerUp SYBR Green Master Mix (Thermo Fisher Scientific), 0.5 μM of each primer and 4 μL of template DNA. qPCR was performed on QuantStudio 3 and 5 Real-Time PCR Systems (Thermo Fisher Scientific) based on the following cycling parameters: 95°C for 3 min, 35 cycles of 95°C for 10 s, 55°C for 10 s for the gp01 and gp38 primers or 57°C for the gp26 primers and 72°C for 15 s. The amplification efficiencies of *gp01, gp26* and *gp38* were validated (Figure S2) and the amount of viral DNA in individual samples at different locations was quantified using their C_T_ values, averaged from three technical replicates.

### Plaque assay

Plaque assay was performed as described previously [22]. Briefly, serially diluted virus preparations, 100 μl each, were pre-mixed with 2 mL of fresh Δarrays/p*gp16* cells (OD_600_ around 0.2) and incubated at 78°C for 30 minutes. Virus-infected culture was mixed with an equal volume of 0.4% Gelzan™ CM (Merck) and spread onto pre-warmed 0.7% Gelzan™ CM/SCV plates. Plates were incubated at 78°C for 2 days, following which the plaques were counted to determine the virus titer.

### Sequencing and statistical analysis

200 mL cell cultures of Δarrays/pcontrol and Δarrays/p*gp16* were grown to an OD_600_ of 0.2 and infected with Δ*gp16*-a. Samples of 15 mL were taken at 1 h, 2 h and 4 h post infection, centrifuged at 6,300x g and cell pellets were washed three times with basic medium salts solution [26]. DNAs were purified as described above in the qPCR section. The extracted nucleic acids were further treated with 1 μL of RNase A solution (#R6148, Sigma Aldrich) for 30 min at 37°C, and purified with DNeasy Blood & amp Tissue Kit (QIAGEN GmbH, Hilden, Germany). Subsequently, 2.5 μL of the purified DNA was amplified by Multiple Displacement Amplification (MDA) with the Genomephi V3 kit (GE Healthcare Life Science, Marlborough, MA, USA) for 30 minutes. Finally, the amplified DNA was purified with a Genomic DNA Clean & amp Concentrator™ kit (Zymo Research, Irvine, CA, USA). The concentration of the MDA amplified and purified DNA was measured by Qubit dsDNA HS Assay Kit (ThermoFisher Scientific, Waltham, MA, USA). Random shotgun libraries were constructed using the Nextera XT kit (Illumina, San Diego, CA, USA) and normalized by AMPure XP beads according to the manufacturer’s protocol. Constructed libraries were sequenced using 2×150 bp paired-end settings on an Illumina NextSeq platform.

To estimate the replication initiation sites, the per-base coverages of the samples from the infected cultures were normalized with that of the virion DNA in 500 bp sliding windows. In short, the raw reads were aligned to the viral genome (SIRV2M_VIII_/Δ*gp16*-a) with Bowtie 2 (v2.4.2) [30]. The per-base coverage was calculated with Samtools [31] (v1.11), and the coverage of the infection samples were divided by that of the virion DNA sample to remove sequencing bias, and the normalized coverage was reported in sliding windows of 500 bp.

### Quantification of the parental and mutant viruses in competition assays

The parental virus and Δ*gp16*-b mutant were mixed at 1:10 before infection. Samples of 1 ml were taken at time points indicated in Results, centrifuged at 6,300 xg and 1 μL of normalized samples were used as template in PCR using primers listed in Table S1. PCR reactions were performed in 20 μl mixtures (Phanta Max Super-Fidelity, Vazyme) as described in Table S5. PCR products were separated in agarose gels of 1.4%. DNA bands in agarose gels were quantified using ImageJ [32]. Relative abundances of each virus at all time points were normalized to the 0 hpi sample (Figure S5).

### Phylogenetic analysis of Rep protein

Homologs of SIRV2 gp16 (Rep) were identified with PSI-BLAST [33] against NCBI protein database. The homologous sequences were retrieved and aligned with Clustal Omega [34] and a phylogenetic tree was constructed using neighbor-joining method. The tree data was visualized and modified using iTOL [35].

## RESULTS

### Design of CRISPR-based genome editing for knocking out conserved genes and deletion of *gp16* from SIRV2

CRISPR based deletion of non-conserved genes in SIRVs has been described earlier [21]. Conserved genes are potentially more important for the basic life cycle of viruses and therefore more difficult to manipulate genetically. Here we developed a modified version, of the previously described deletion technique, to knock out genes conserved among SIRVs, involving two different host strains (Figure 1A). The first strain, referred as the deletion strain, carries a plasmid containing a CRISPR subtype I-A mini-array, necessary for CRISPR-mediated target DNA cleavage, and the homologous recombination arms, derived from viral sequences flanking the CRISPR target site. Upon virus infection of the deletion strain, we expect to see a mixture of the parental virus and the mutant virus, with the former persisting as a result of the fitness advantage provided by the target gene and the latter arising due to the CRISPR-Cas selection pressure. The second strain, referred as the purification strain, carries a plasmid encoding a mutated version of the target gene along with a CRISPR subtype I-A mini-array. The crRNA transcribed from this mini-array is complementary to the target gene present in the parental virus but incapable of mediating targeting of the mutated version carried on the second plasmid. Plasmid-borne gene complementation enables the mutant virus propagate at the same rate as the parental virus, but only the parental virus is targeted by the CRISPR-Cas system (Figure 1A). Based on this principle, pure mutant viruses can be obtained by infecting the purification strain with the virus mixture obtained from the infected deletion strain (Figure 1A).

SIRV2 gp16 protein belongs to the replication initiator (Rep) superfamily [12]. To study the *in vivo* function of the Rep protein, we intended to knock out *gp16* from the SIRV2M_VII_ mutant which lacks the accessory genes present among *Rudiviridae* and does not show any significant fitness disadvantage compared to the wild-type SIRV2 virus [21].

Supernatants from the infected deletion strain were checked by PCR, a DNA band indicative of *gp16* deletion (237 bp in length) was detected after one round of infection and the parental virus was still present (568 bp in length) (Figure 1B; Lane 1). The virus mixture was transferred into the culture of purification strain. After two rounds of passages the pure *gp16*-lacking viral mutant was obtained (Figure 1B; Lane 2). In accordance with the nomenclature applied earlier [21], the mutant virus was named SIRV2M_VIII_, referred to as Δ*gp16*-a (Table S2). Mutant virus purity was further confirmed by PCR using primers annealing to *gp16* sequence (Figure 1B, right panel).

### Impact of Rep protein on viral yield

First, we assessed the effect of Rep on virus propagation. *S. islandicus* LAL14/1 Δarrays strain was transformed with a plasmid carrying SIRV2 *gp16* with its native promoter, and the resultant transformant was referred as Δarrays/p*gp16* (Table S3). *S. islandicus* LAL14/1 Δarrays/pcontrol (empty vector) and the complementation strain Δarrays/p*gp16* were infected with Δ*gp16*-a at an MOI of 0.001. As controls the strains were also infected with the parental virus SIRV2M_VII_ (WT).

Complemented by a plasmid-borne *gp16*, the mutant virus inhibited host growth similarly as the parental virus; however, in the absence of *gp16* the host growth was much less affected by the infection of the mutant virus (Figure 2A). Accordingly, the absence of *gp16* decreased the yield of the mutant virus 4-6 fold at 13 hpi, the end of a complete SIRV2 life cycle [10, 16], in comparison to the parental virus (Figure 2B). This suggests that, although not essential, Rep enhances SIRV2 propagation in the Δarrays strain.

**Figure 2.**
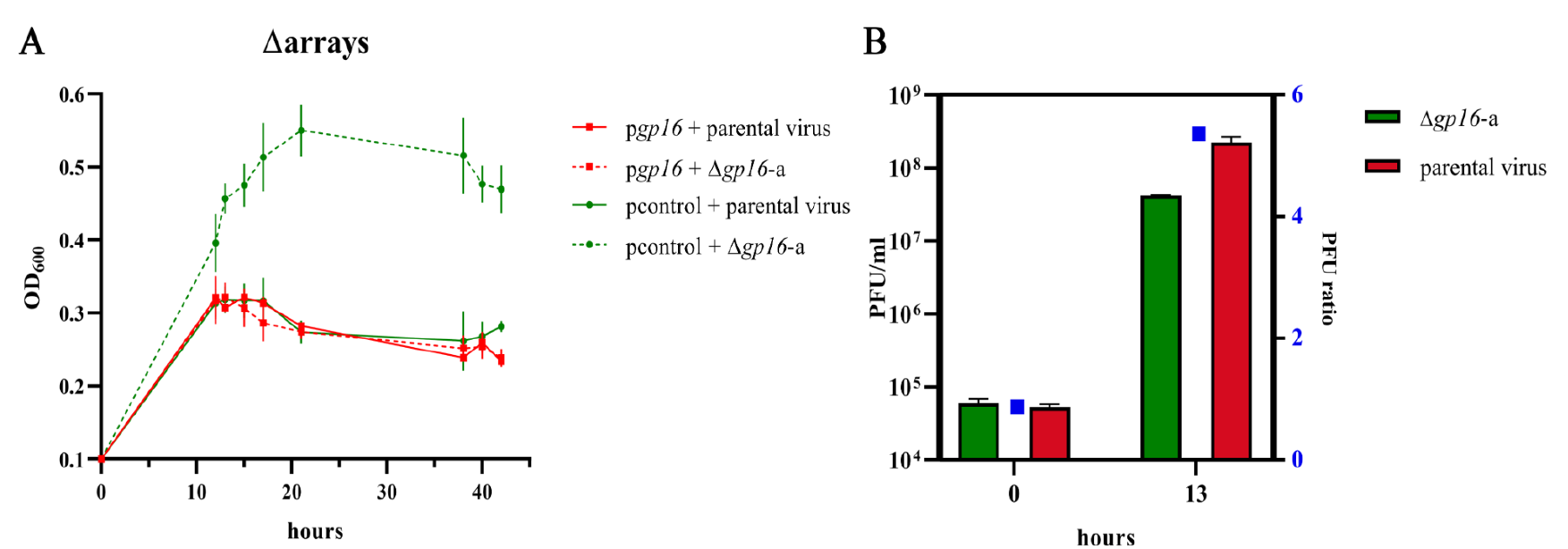
Physiological effect of gp16 on virus propagation. (A) Growth curves of *S. islandicus* LAL14/1 Δarrays strain carrying the empty plasmid (pcontrol) or the complementation plasmid (p*gp16*) infected with Δ*gp16*-a or the parental virus at an MOI of 0.001. (B) Extracellular virus titers of *S. islandicus* LAL14/1 Δarrays infected with the parental virus (red) or Δ*gp16*-a (green) (shown earlier) at 0 and 13 hpi. PFU, plaque forming units. Results shown here are a mean of three biological replicates and error bars indicate the corresponding standard deviations (SD).

### Rep initiates replication at SIRV2 genomic termini

The Rep protein was shown *in vitro* to make a nick at four sites of the SIRV2 genomic termini sharing a 4-nt motif [12], and more recently, multiple replication initiations were observed from the termini of single parental SIRV2 genomes [13]. We therefore hypothesized that the increase in viral yield conferred by the Rep protein (Figure 2B) was due to a more efficient replication. Therefore, we decided to quantify the intracellular SIRV2 DNA in the presence and absence of Rep. The aforementioned strains, Δarrays/pcontrol and Δarrays/p*gp16*, were infected with Δ*gp16*-a at an MOI of 0.1 and sampled between 0 and 8 hpi. Quantitative PCR was performed on the extracted total genomic DNA to quantify intracellular viral DNA at three different genomic locations using primers specific to the terminal (*gp01*), quarter (*gp26*) or central (*gp38*) region of Δ*gp16*-a.

First, we quantified extracellular virions at multiple time points within a single virus life cycle. Correlating to results shown earlier (Figure 2B) there was a 2-8 fold increase in virus yield in the presence of Rep (Figure 3A) starting at 4 hpi, when the earliest virus release was observed [10]. Intracellular viral DNA counts measured at the genomic termini were significantly higher in the presence of Rep than in the absence of Rep and the difference peaked at 2 hpi (about 6 fold) (Figure 3B). The subsequent drop in difference at 4 hpi (2 fold) (Figure 3 B) coincided with the initial virion release (Figure 3A). The second peak, at 6 hpi (3 fold), is probably derived from a second round of infection because we expect less than 10% of the cells to be infected in the first round given the low MOI of 0.1 and virions released at around 4 hpi [10, 16] can infect the remaining 90% cells (Figure 3 A-B). On the other hand, intracellular viral DNA counts measured at the non-terminal regions were affected only minimally by Rep (Figure 3 C-D). These results indicate that Rep specifically increases DNA counts at the viral genomic termini, suggestive of Rep-dependent replication initiation at the termini. Importantly, although viral DNA replication still prevails, it is significantly affected by the absence of the Rep protein.

**Figure 3.**
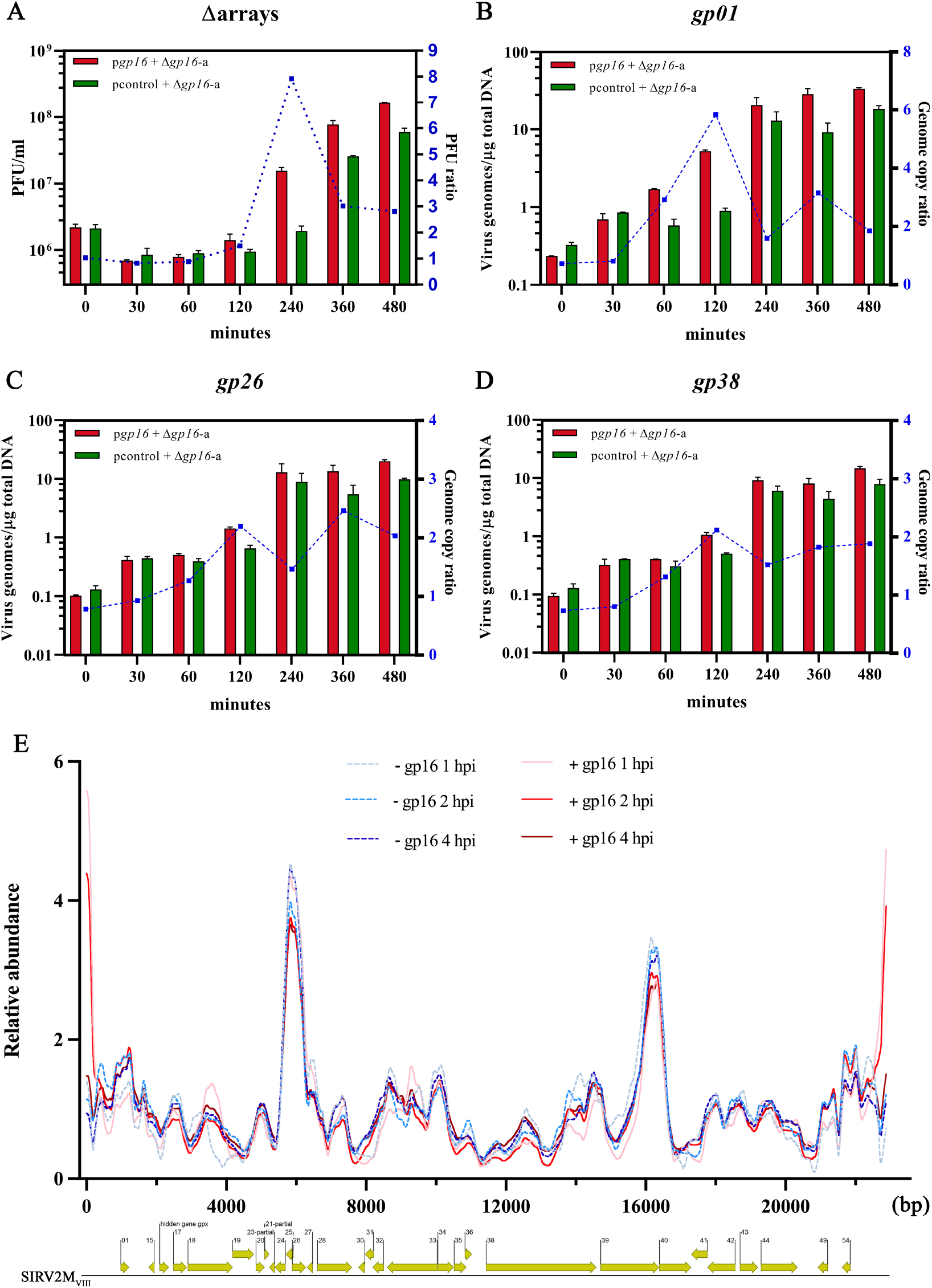
Rep initiates replication from the viral genomic termini. (A) Virus titers. *S. islandicus* LAL14/1 Δarrays/p*gp16* and Δarrays/pcontrol were infected with Δ*gp16*-a at an MOI of 0.1. Samples were collected at multiple time points (min) post infection and extracellular virus titers were measured by plaque assay. PFU ratio: ratio of virus titer between gp16-containing and gp16-null cultures. (B-D) Intracellular viral genome copy as measured by qPCR using primers specific for the viral genomic end region (*gp01*), quarter region (*gp26*) and central region (*gp38*). Genome copy ratio, viral genome copy ratio between gp16-containing and gp16-null cultures. (E) Relative abundance of sequencing reads across the viral genome of Δ*gp16*-a (bp). The genome map of Δ*gp16*-a is presented below to show the corresponding positions of the genes, and the gp (gene product) numbers correspond to those of SIRV2. Data were generated from *gp16*-containing cultures (red lines) and *gp16*-null cultures (blue dashed lines) sampled 1, 2 and 4 hpi. Results shown here are a mean of duplicate biological replicates and error bars indicate the corresponding SD.

To obtain more evidence for the function of Rep in SIRV2 replication initiation, we infected Δarrays cells with the Δ*gp16*-a viral mutant in the presence or absence of a plasmid-borne *gp16*, extracted total DNA from cells sampled at 1, 2 and 4 hpi and sequenced the DNA using Illumina sequencing. SIRV2 virion DNA was sequenced as a non-replicating control. Sequence reads of the six samples were normalized using reads generated from SIRV2 virion DNA and plotted against Δ*gp16*-a genome.

In the presence of Rep, sequence coverage is up to 5 fold higher at genomic termini than that in the absence of Rep, further supporting the role of Rep in replication initiation at the ends. Interestingly, two internal sequencing peaks, located around *gp26* and *gp39*, respectively, were observed in both types of samples (Figure 3E). We hypothesize that these two sites serve as internal replication initiation origins, contributing to a basal level of virus replication.

Taken together, these results indicate that the Rep protein is responsible for replication initiation at the termini and there exists alternative origin(s) of replication which become the single source of replication initiation in the absence of the Rep protein.

### Rep protein provides virus with a selective advantage in host-virus arms race

Despite the importance of Rep in replication initiation at SIRV2 genomic termini (Figure. 3E), its absence led to only about 8 fold drop in virus yield in Δarrays cells (Figure 3A) which carries no CRISPR immunity. We reasoned the phenotype could be different in a more natural setting with the presence of host CRISPR-Cas immunity and viral anti-CRISPRs, therefore we set out to compare the infectivity between *gp16* positive and negative viral clones in wild-type LAL14/1. With anti-CRISPR proteins [17, 18] the wild-type SIRV2 exhibits a high infectivity in the wild-type LAL14/1 host despite the presence of one type I-A, one I-D and two III-B CRISPR-Cas systems in the latter [16]. CRISPR-based genome editing of the wild-type SIRV2 is, however, not possible in LAL14/1 whose CRISPR-Cas systems are all inhibited by SIRV2 Acrs. As such, *gp16* was deleted from the virus SIRV2M_II_*gp02* which does not carry the Acr gene *acrIIIB1* (Figure S3, Table S2). The infectivity of the resultant Δ*gp16*-b was then compared with that of the parental virus after infecting the wild-type *S. islandicus* LAL14/1 strain carrying a plasmid-borne *acrIIIB* (referred as LAL14/1-p*acrIIIB2)*. As a control, the Δarrays/pcontrol strain was also infected with the two viruses. For both viruses, the initially selected MOI was 10^−5^, and the host growth and virus titers were monitored over a period of 72 hours.

While both viruses caused growth retardation of the infected Δarrays/pcontrol strain, and so did the parental virus in the infected wild-type host, Δ*gp16*-b infection did not impair the growth of the wild-type host (Figure 4A). In correlation with this, the titer of Δ*gp16*-b in the wild-type host remained constantly lower than that of the parental virus reaching a maximum difference of ∼180 fold at 60 hpi (Figure 4B and 4D). In contrast, in Δarrays/pcontrol strain, the presence of *gp16* resulted in a maximum of 20 fold difference at 12 hpi and showed no significant difference at any other time points (Figure 4C and 4D). The ratio of virus titer between the parental virus and Δ*gp16*-b in both hosts is depicted in Figure 4D.

**Figure 4.**
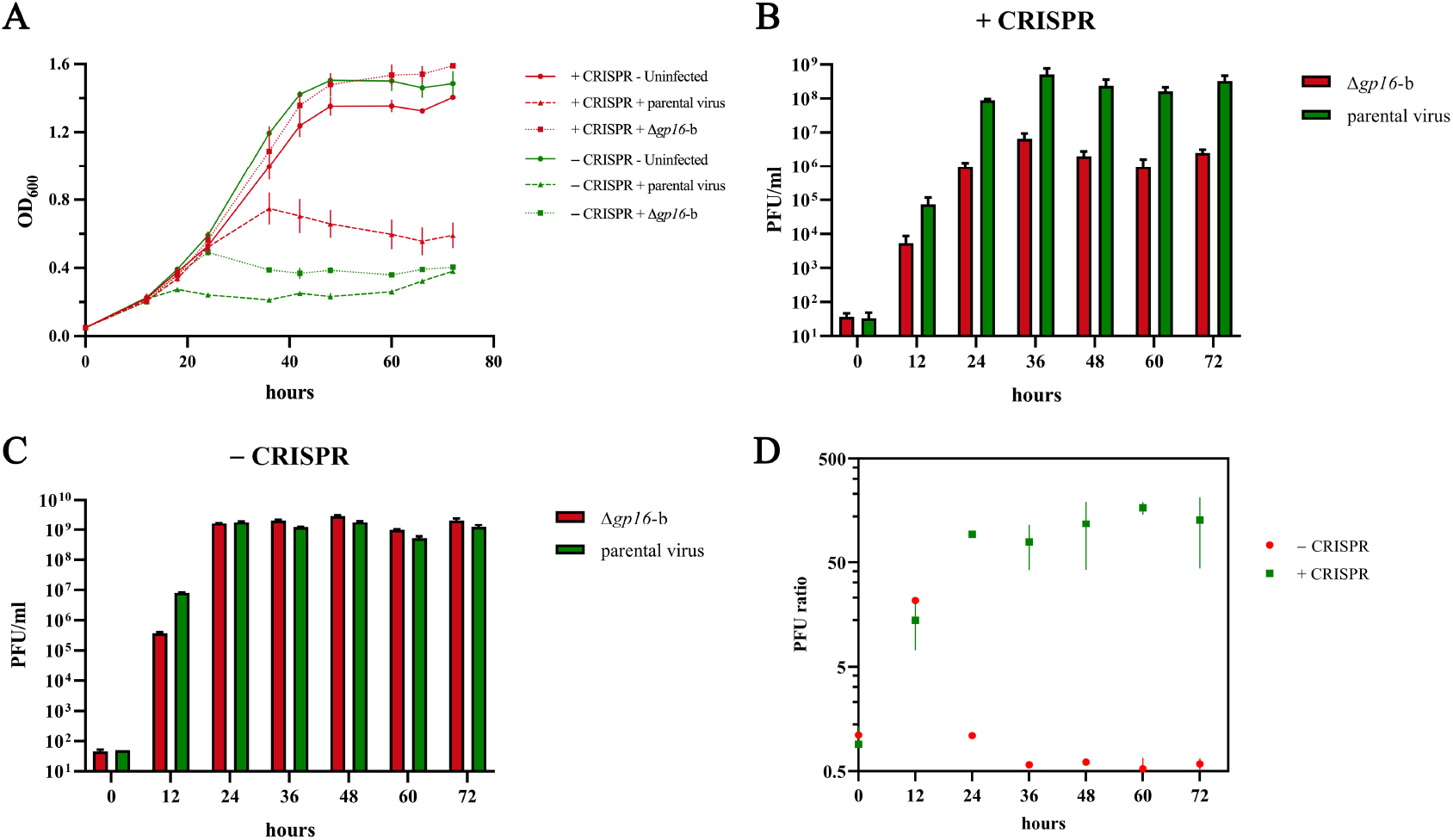
Rep protein provides selective advantage to virus propagation in the presence of CRISPR-Cas immunity. (A) Growth curves of *S. islandicus* LAL14/1 wild-type (+ CRISPR) and Δarrays (-CRISPR) infected with Δ*gp16*-b or its parental virus (SIRV2M_II_*gp02*) at MOI = 10^−5^ measured over 72 hours. (B) Extracellular virus titers of Δ*gp16*-b (red) and the parental virus (green) measured at 12-hour intervals in *S. islandicus* LAL14/1 which carries functional CRISPR-Cas systems. (C) Extracellular virus titers of Δ*gp16*-b (red) and the parental virus (green) measured at 12-hour intervals in Δarrays cells which does not carry a functional CRISPR-Cas system. (D) Ratio of extracellular virus titer between the parental virus and Δ*gp16*-b (as measured in B and C) in *S. islandicus* LAL14/1 wild-type strain (+ CRISPR) and Δarrays (-CRISPR) over 72 hours post infection. Results shown here are a mean of three biological replicates and error bars indicate the corresponding SD. PFU, plaque forming unit.

In addition, we also quantified the virion yield of the parental virus and Δ*gp16*-b in the wild-type host at different MOIs (10^−4^∼10^−7^) at 72 hpi, a time window corresponding to 5 – 6 SIRV2 life cycles [10, 16]. The parental virus retards growth of the wild-type host at all MOIs except at an MOI of 10^−7^, however only a mild growth retardation was observed even at the highest titer of infection for the mutant virus (Figure S4A). At all MOIs, the parental virus showed a higher yield than Δ*gp16*-b, and, a 1000-fold difference in virus yield was observed at the MOI of 10^−6^ (Figure S4B). Taken together, these results show that in the presence of CRISPR-Cas immunity, the lack of Rep led to a more profound drop of infectivity than in the host without CRISPR-Cas immunity.

To better understand the importance of Rep for SIRV2 propagation in a competitive environment, we co-infected LAL14/1-p*acrIIIB2*, Δarrays/pcontrol and Δarrays/p*gp16* with a mixture of the parental virus and Δ*gp16*-b at a ratio of 1:10. Viruses in the culture supernatants were diluted into fresh cultures every 48 hours for up to 192 hours and sampled every 24 hours for semi-quantitative comparison using PCR. Supernatants were normalized for total virus quantity with a primer pair specific for a region conserved in both viruses. Virus specific primers (Table S1) were used to differentiate between the parental virus and Δ*gp16*-b. In *S. islandicus* LAL14/1 wild-type strain, Δ*gp16*-b was quickly outcompeted by the parental virus, as shown by the barely visible PCR band at 72 hpi (Figure 5A, left panel; Figure S5A). In contrast, in the absence of CRISPR-Cas immunity the mutant virus persisted for a longer period, until 168 hpi, when the parental virus eventually outcompeted the mutant virus (Figure 5A, middle panel; Figure S5B). In the presence of a plasmid-borne *gp16* (Δarrays/p*gp16*) the Δ*gp16*-b mutant propagated at the same rate as the parental virus (Figure 5A, right panel; Figure S5C). Next we wanted to examine the role of the Rep protein in the presence of individual CRISPR-Cas systems. Similar to the results observed with the wild-type host, Δ*gp16*-b virus was rapidly outcompeted by the parental virus, independent of the CRISPR-Cas subtype encoded by the host (Figure 5B, Figure S5D, S5E and S5F). The rate at which the mutant virus was outcompeted was similar in hosts carrying type I CRISPR-Cas systems (subtypes I-A and I-D) but lower in the host carrying the subtype III-B system, which could be attributed to the mechanistic differences associated with the different CRISPR-Cas targeting activities. Taken together, these results indicate that rudiviruses encoding gp16 have a selective advantage in virus-virus conflicts, especially in the presence of CRISPR-Acr warfare these differences are substantially magnified.

**Figure 5.**
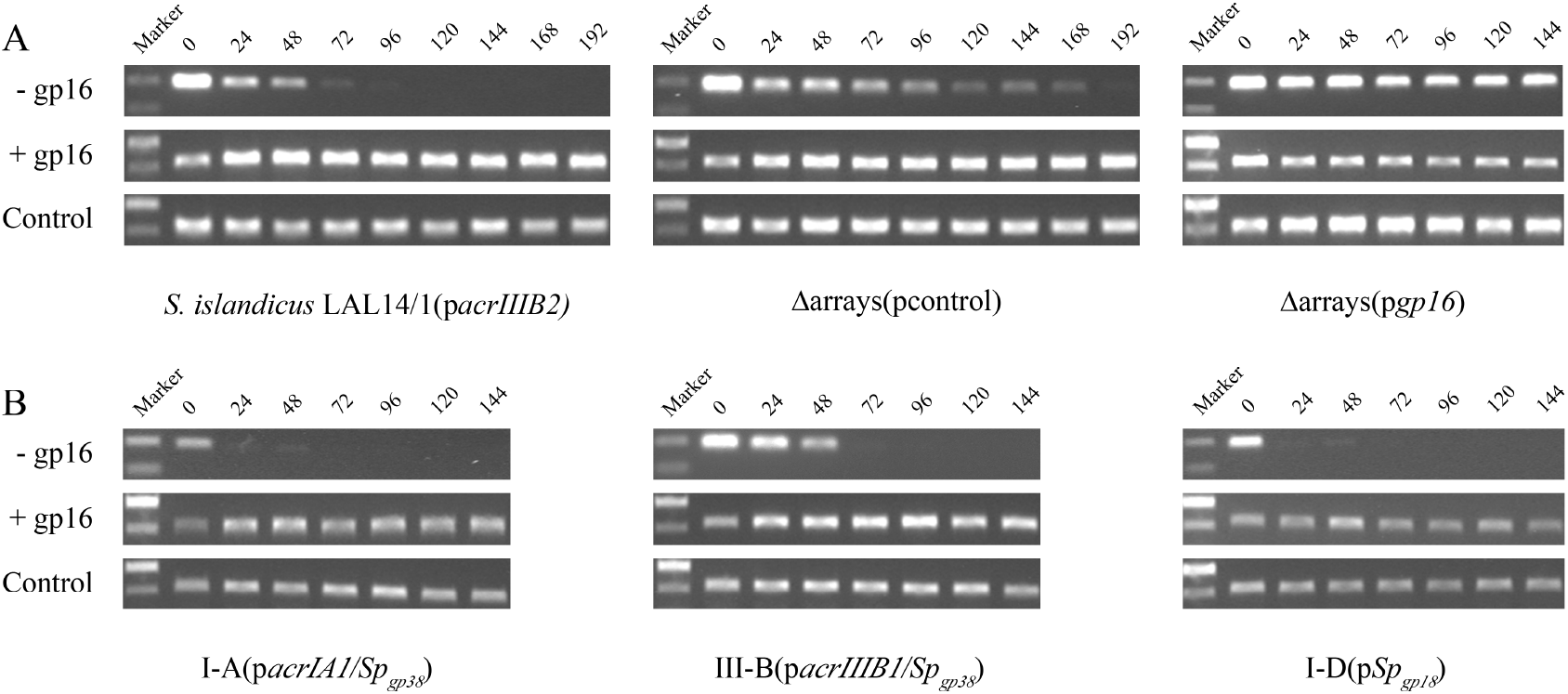
Virus competition upon co-infection of different *S. islandicus* LAL14/1 strains. (A) PCR amplification of the parental virus (+ gp16) and Δ*gp16*-b (-gp16) present in the supernatants of co-infected *S. islandicus* LAL14/1 carrying CRISPR-Cas systems (Left), Δarrays/pcontrol (Middle) and Δarrays/p*gp16* (Right). (B) Same as (A) but *S. islandicus* LAL14/1 strains carrying a single CRISPR-Cas system, either I-A (Left), III-B (Middle) or I-D (Right), were used. Sampling times (hpi) are indicated on top. Control, PCR products using primers specific to *gp26*-coding sequence. All primers are listed in Table S1. Marker, DNA size marker.

### Rep protein is conserved in Rudiviridae

PSI-BLAST identified 22 homologs of gp16 protein, with common characteristic being the conserved HUH domain and a tyrosine residue typical of HUH endonuclease superfamily Rep proteins (Figure S6). Phylogenetic analysis of the Rep protein homologs resulted in grouping of the proteins from the same geographical location of isolation. Interestingly, all the 22 homologs were specifically restricted to the Rudiviridae family and absent in the other family of the Ligamenvirales order, Lipothrixviridae (Figure 6). Although both families have linear dsDNA genomes with inverted terminal repeats (ITRs) their replication mechanisms have been shown to be very different [13, 36]. The termini of lipothrixviral genomes are not covalently linked and gp16-mediated rolling circle replication is therefore not relevant.

**Figure 6:**
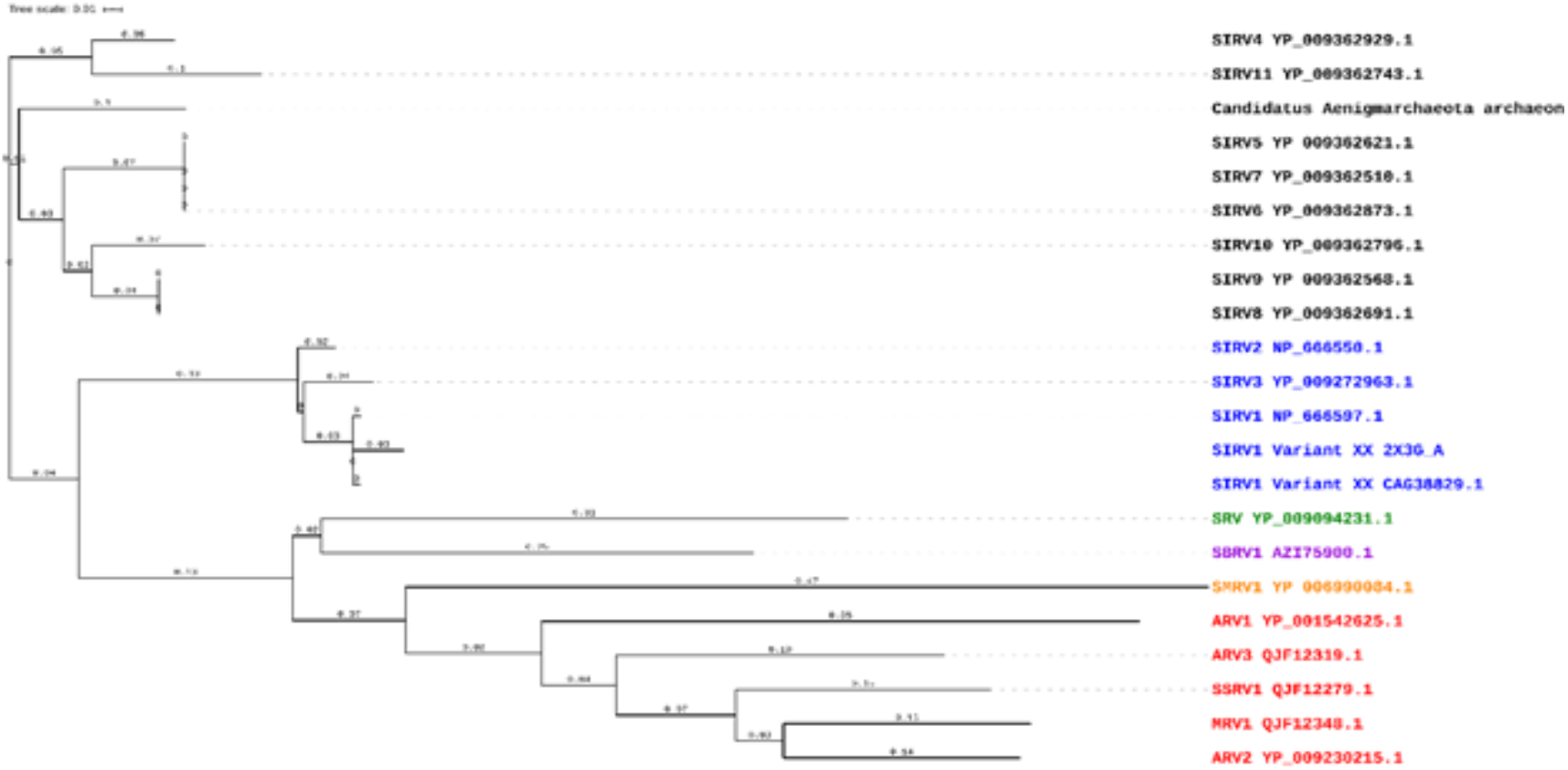
Rep protein homologs are absolutely conserved in *Rudiviridae*. Phylogeny of Rep protein in members of *Rudiviridae*. Construction of the phylogenetic tree is described in materials and methods. The GenBank identifier and gene names are indicated for each sequence. Geographic origins of the corresponding viruses are color-coded, Black, USA; blue, Iceland; Red, Italy; Green, Portugal; Purple, Japan; Yellow, Mexico.

## Discussion

Whereas earlier structural [37] and *in vitro* enzymatic data [12] indicated a replication initiator role for SIRV2 gp16, two independent transcriptomic studies [10, 11] showed extremely low expression of *gp16* in two different *Sulfolobus* hosts. The absolute number of reads per kilobase of transcript for *gp16* ranged from 0 to 250 within 9 hpi which is one to three orders of magnitude lower than those of other SIRV2 genes [11]. This raised doubts about its role in replication. We demonstrate in this study that gp16 is indeed a replication initiator as evidenced by the disappearance of replication initiation from the genomic termini of *gp16* knockout mutant. The 4-6 fold decrease in viral yield, resulting from the deletion of gp16, demonstrates that gp16 is responsible for at least 80% of SIRV2 replication initiation under the tested conditions (Figure 2B).

Apart from the terminal replication origins, two replication origins appear to exist internally, one around SIRV2 *gp26*, the other around *gp39* (Figure 3E). Multiple replication origins were also reported for viruses infecting bacteria and eukaryotes, as exemplified by phage T4 and Herpes Simplex Virus (HSV) [38, 39]. Similarly, deletion of some of the origins was not completely detrimental for the viruses, but only reduced the viral yield. It’s important to note that the viral genomes are much smaller than bacterial chromosomes which encode only one replication origin [40]. Moreover, several viruses were reported to employ more than one replication modes, e.g. origin-mediated and recombination-dependent DNA synthesis in phage T4 [41]; theta and sigma replication modes of phage lambda [38, 42]; strand-displacement and strand-coupled replication modes of SIRV2 [13]. While it was assumed that the multiple replication origins and replication modes might aid in increasing viral fitness under different environmental stress conditions [38], this study provides the first experimental evidence for the selective advantage of a replication initiator protein, especially in the presence of host CRISPR-Cas immunity.

Intriguingly, despite being unessential for viral propagation (Figure 2A and 2B), *rep* is absolutely conserved with one copy present in each of the known Rudiviridae members isolated from all over the world (Figure 6). Given the ubiquitous presence of CRISPR-Cas systems in their Sulfolobales hosts, e.g. *Sulfolobus, Acidianus, Stygiolobus* [43], and the fact that Rep provides a selective advantage in the presence of host CRISPR-Cas immunity, it’s very likely that the host CRISPR-Cas immunity has driven the evolution of Rudiviridae replication machinery which resulted in the absolute conservation of Rep.

## Supplementary figures

**Figure S1.**
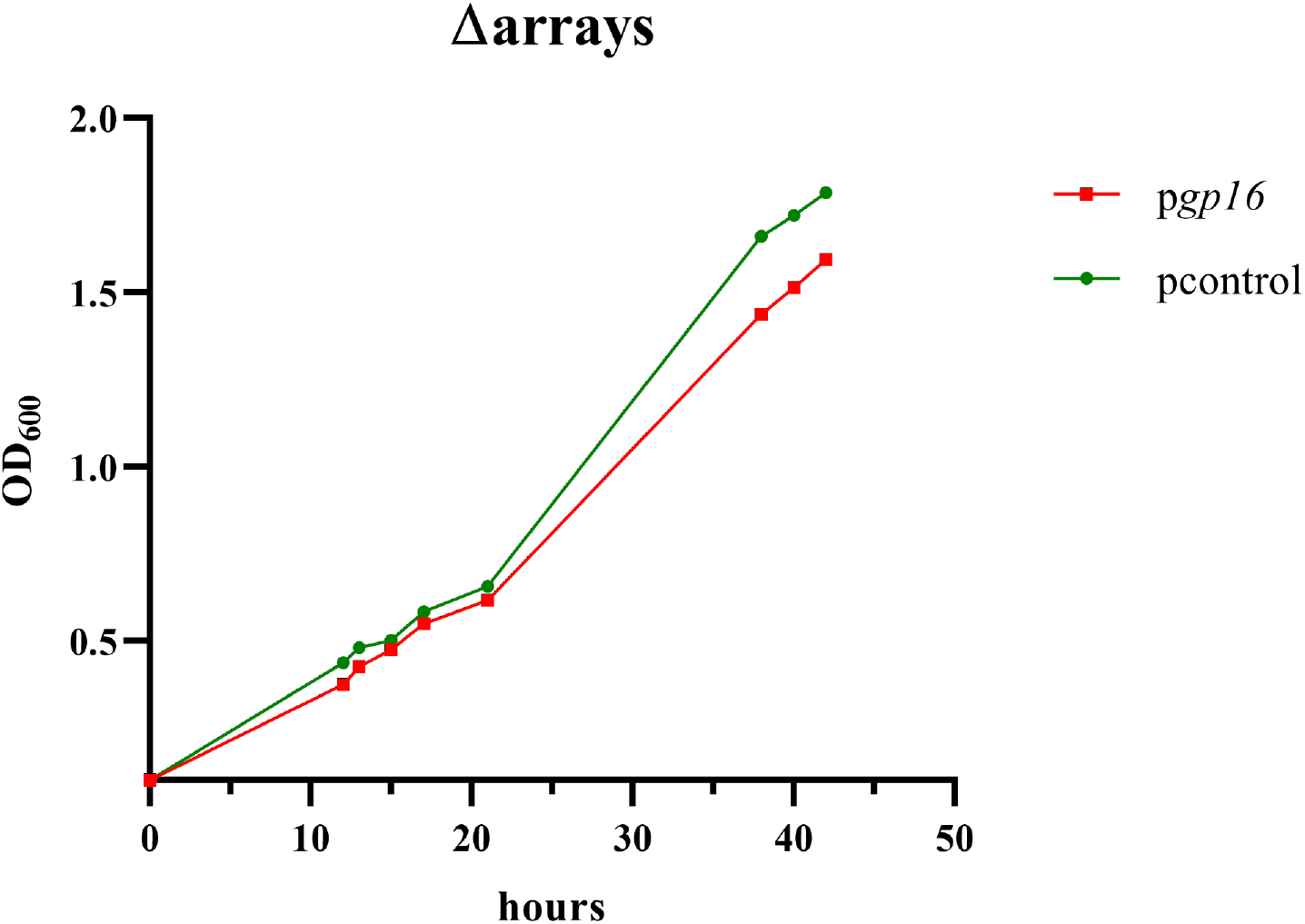
Growth Curves of uninfected cultures of Δarrays/p*gp16* (red) and Δarrays/pcontrol (green).

**Figure S2.**
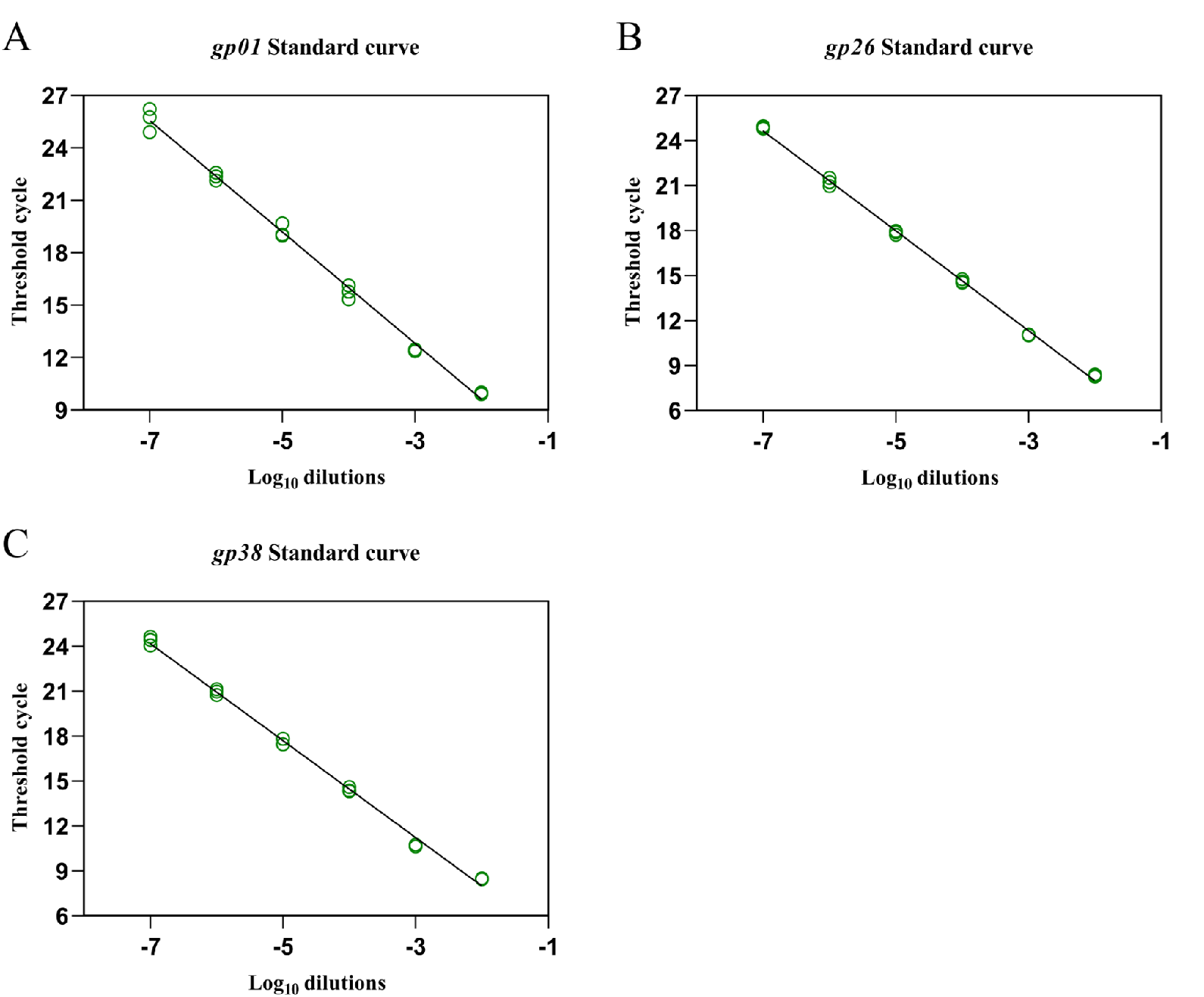
Determination of qPCR efficiencies for primer sets matching *gp01, gp26* and *gp38*. The standard curve was constructed based on SIRV2 virion DNA. The qPCR efficiencies for the standard curves of *gp01, gp26* and *gp38* are 106%, 100% and 104% respectively.

**Figure S3.**
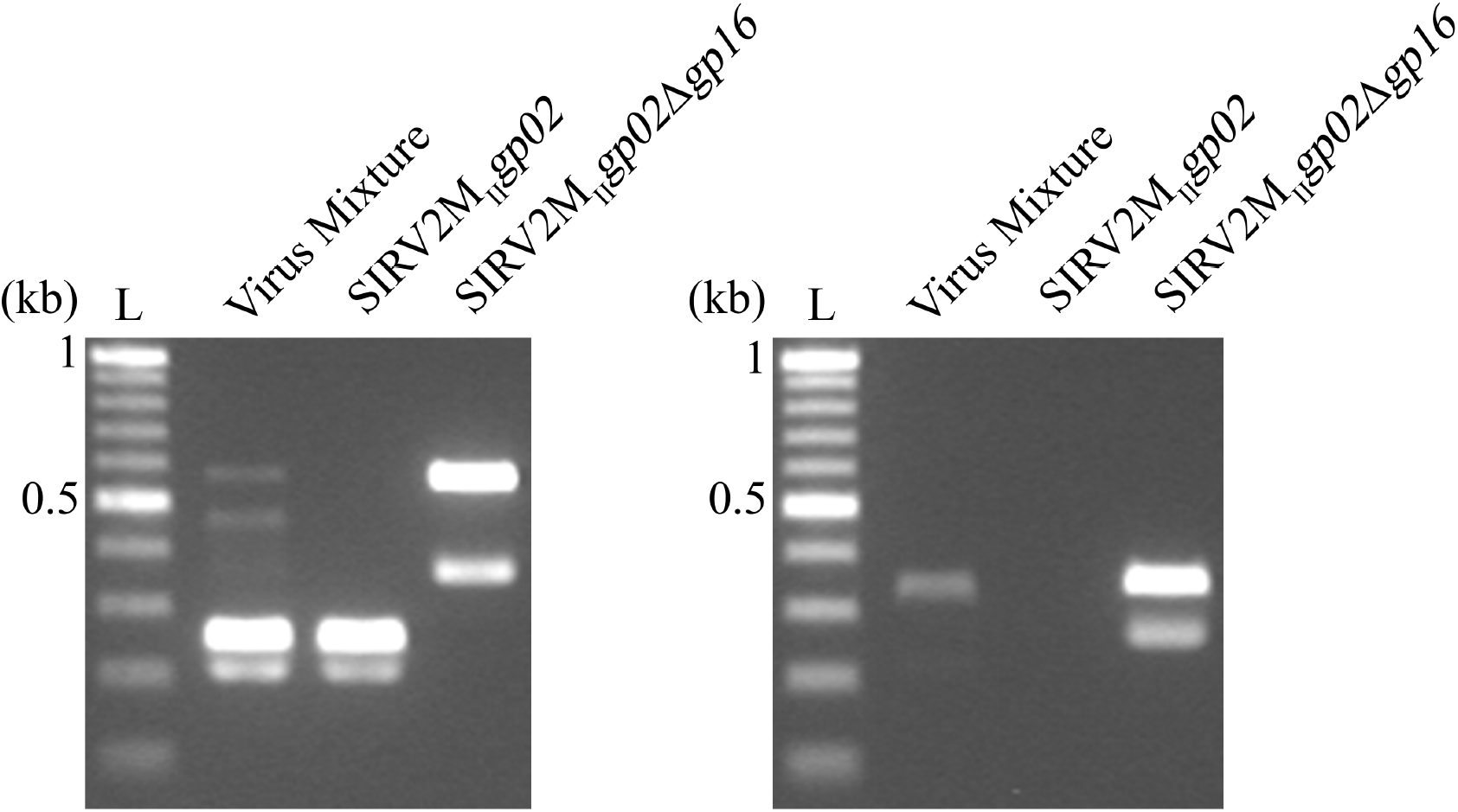
Construction of gp16 deletion mutant, SIRV2M_II_*gp02*Δ*gp16*(Δ*gp16*-b). Left panel, Lane 1 shows a virus mixture containing both mutant and parental viruses, Lane 2 shows purified mutant virus obtained after passage through the purification strain. Right panel Lane 1 and Lane 2, shows virus purity determined with primers specific for the deleted region. Samples were identical to those in the left panel. Lane 3 in both panels represents the parental virus (SIRV2M_II_*gp02*). Primers used in these experiments are listed in Table S1.

**Figure S4.**
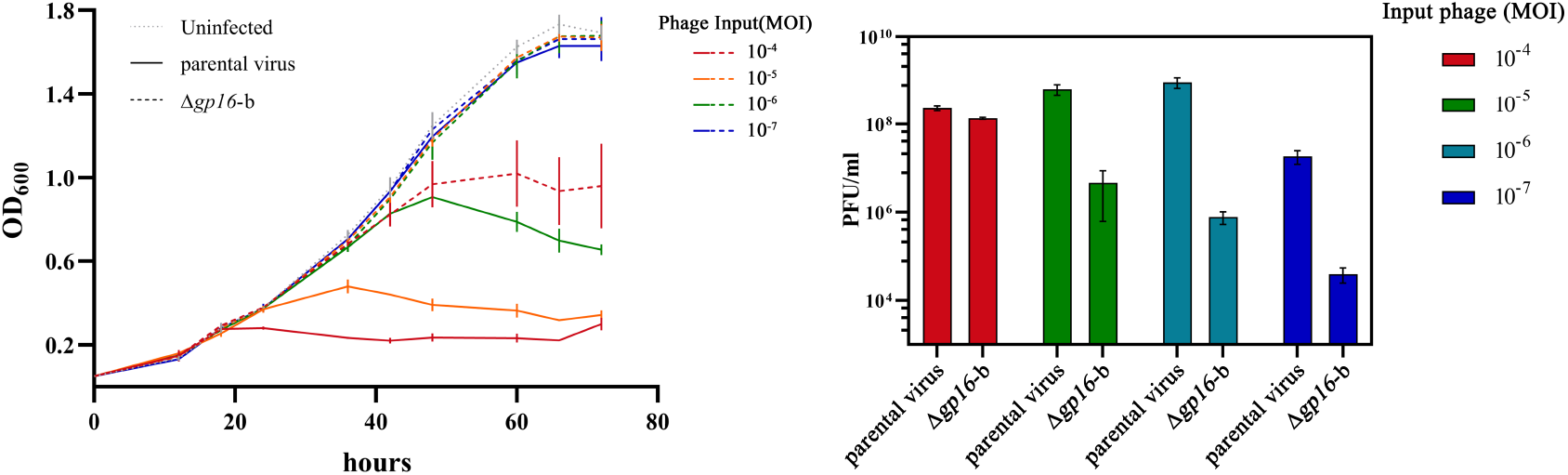
Selective advantage conferred by Rep protein at different MOIs. Left panel, growth curves of *S. islandicus* LAL14/1 infected with the parental virus and Δ*gp16*-b at different MOIs (10^−4^ – 10^−7^). Right panel, extracellular virus titers from the same cultures at 72 hpi.

**Figure S5.**
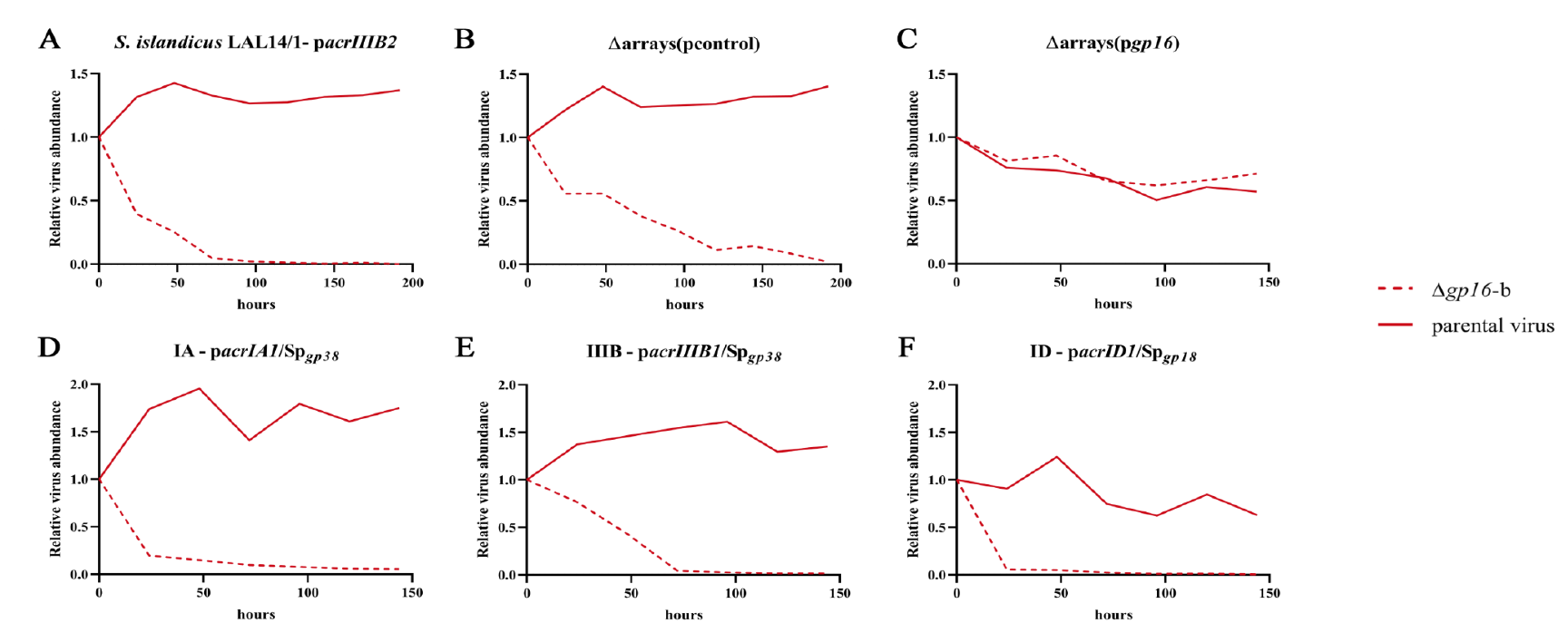
Relative virus abundance. The PCR bands from Figure 5 were quantified using the ImageJ analysis tool. Band intensity at 0 h was set arbitrarily as 1. The abundance of the parental virus is shown as straight lines and that of Δ*gp16*-b as dashed lines.

**Figure S6.**
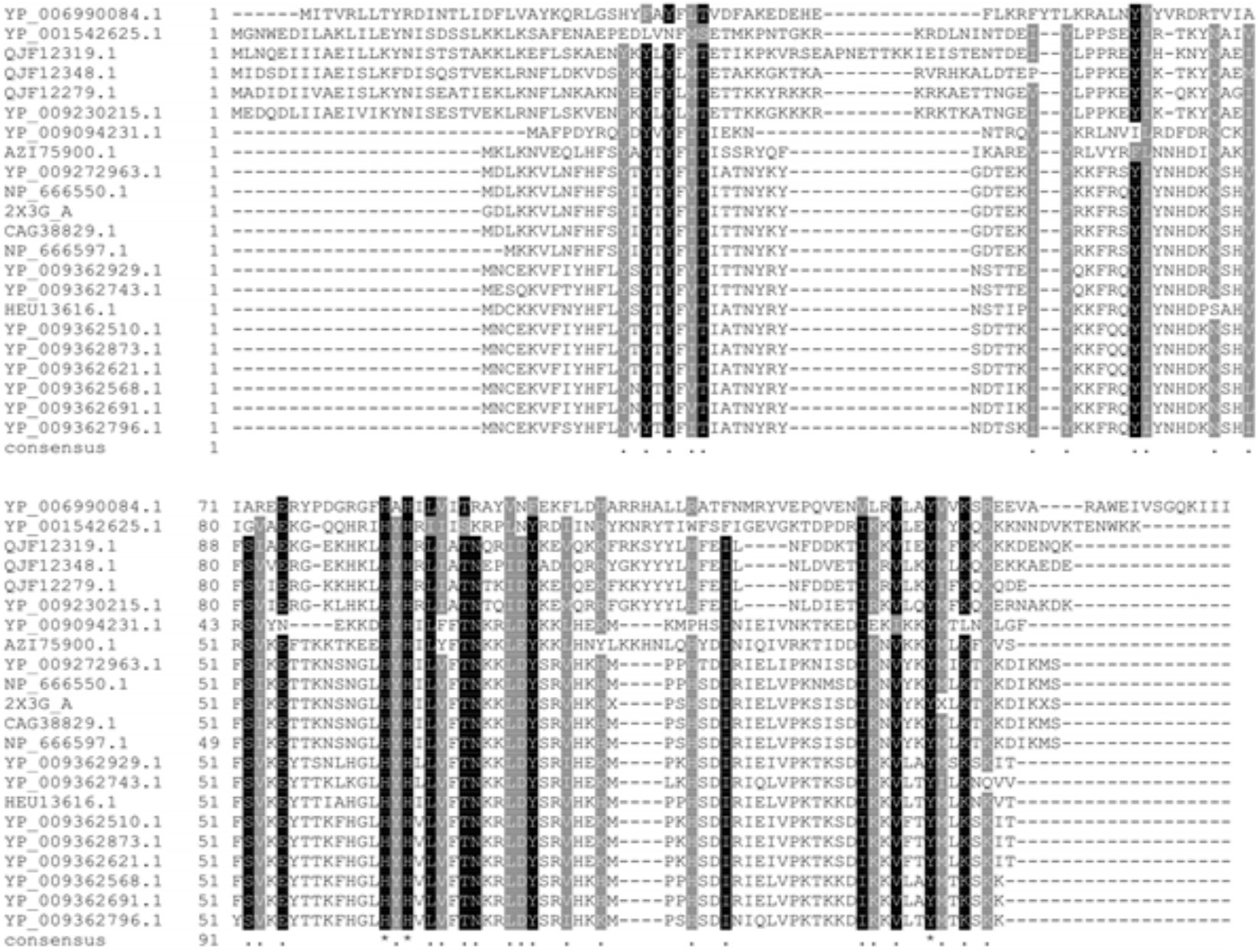
Multiple sequence alignment of Rep protein homologs. The conserved residues are highlighted in black, including the Tyrosine (Y) and HUH motifs.

**Table S1.**
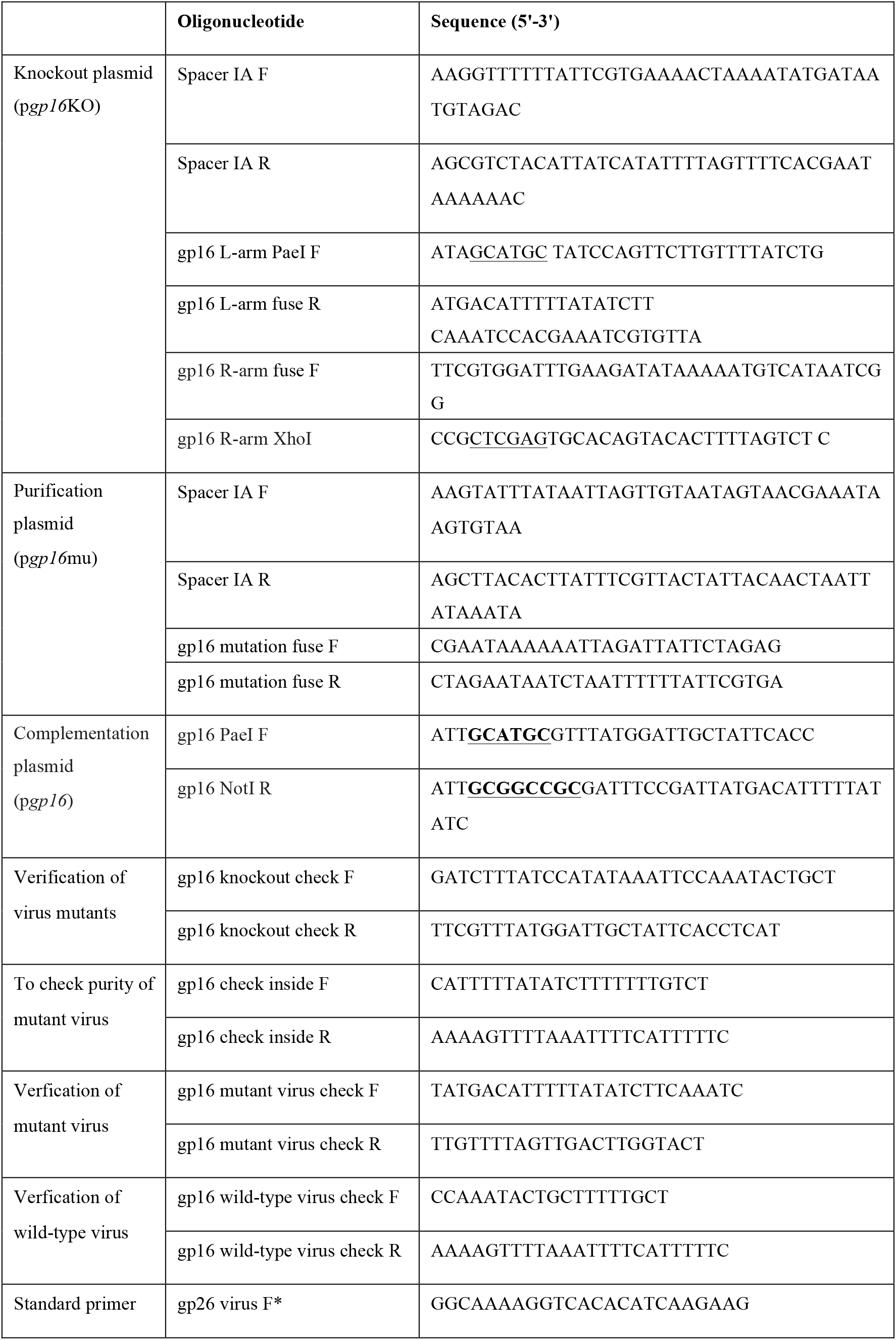
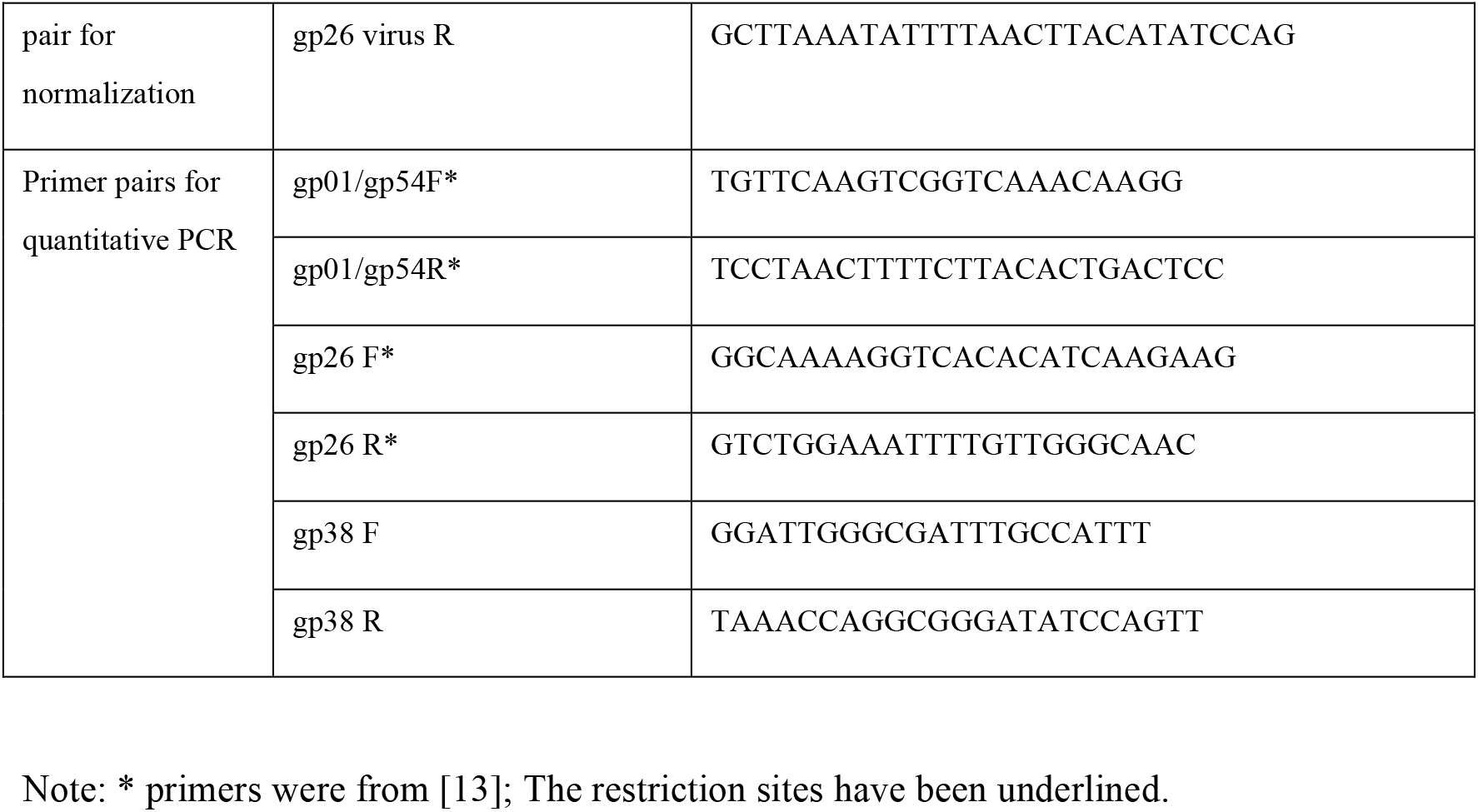
Primer pairs used in this study.

**Table S2.** Viruses.

| Virus | Description | Reference |
| --- | --- | --- |
| SIRV2M<sub>VII</sub> | a SIRV2 mutant with deletions of 25 genes serving as parental virus | [21] |
| SIRV2M<sub>VIII</sub>(Δgp16-a) | mutant virus with deletion of gp16 from SIRV2M<sub>VII</sub> | This work |
| SIRV2M<sub>II</sub>gp02 | a SIRV2 mutant lacking 11 genes and *acrIIIB1*, carrying *acrID1* | [18] |
| SIRV2M<sub>II</sub>gp02Δgp16(Δgp16-b) | mutant virus with deletion of gp16 from SIRV2M<sub>II</sub>gp02 | This work |

**Table S3.**
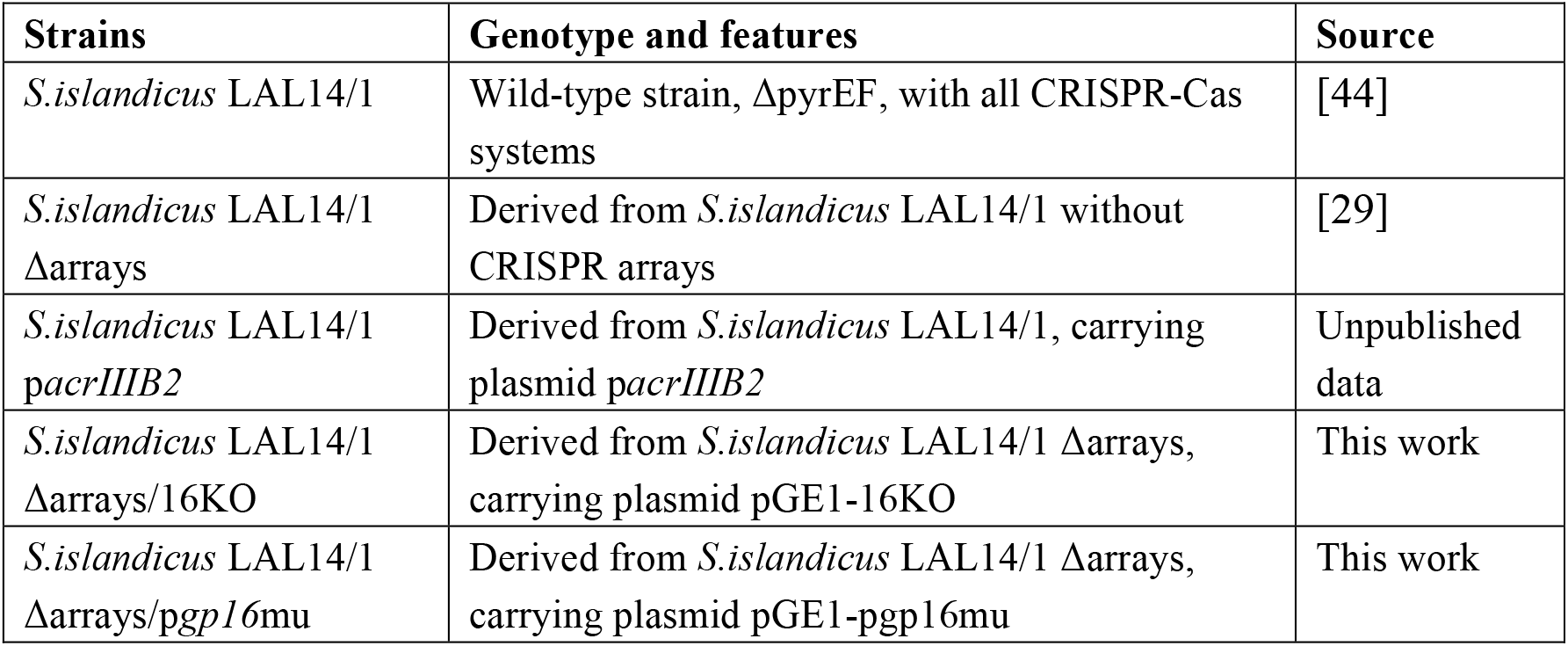
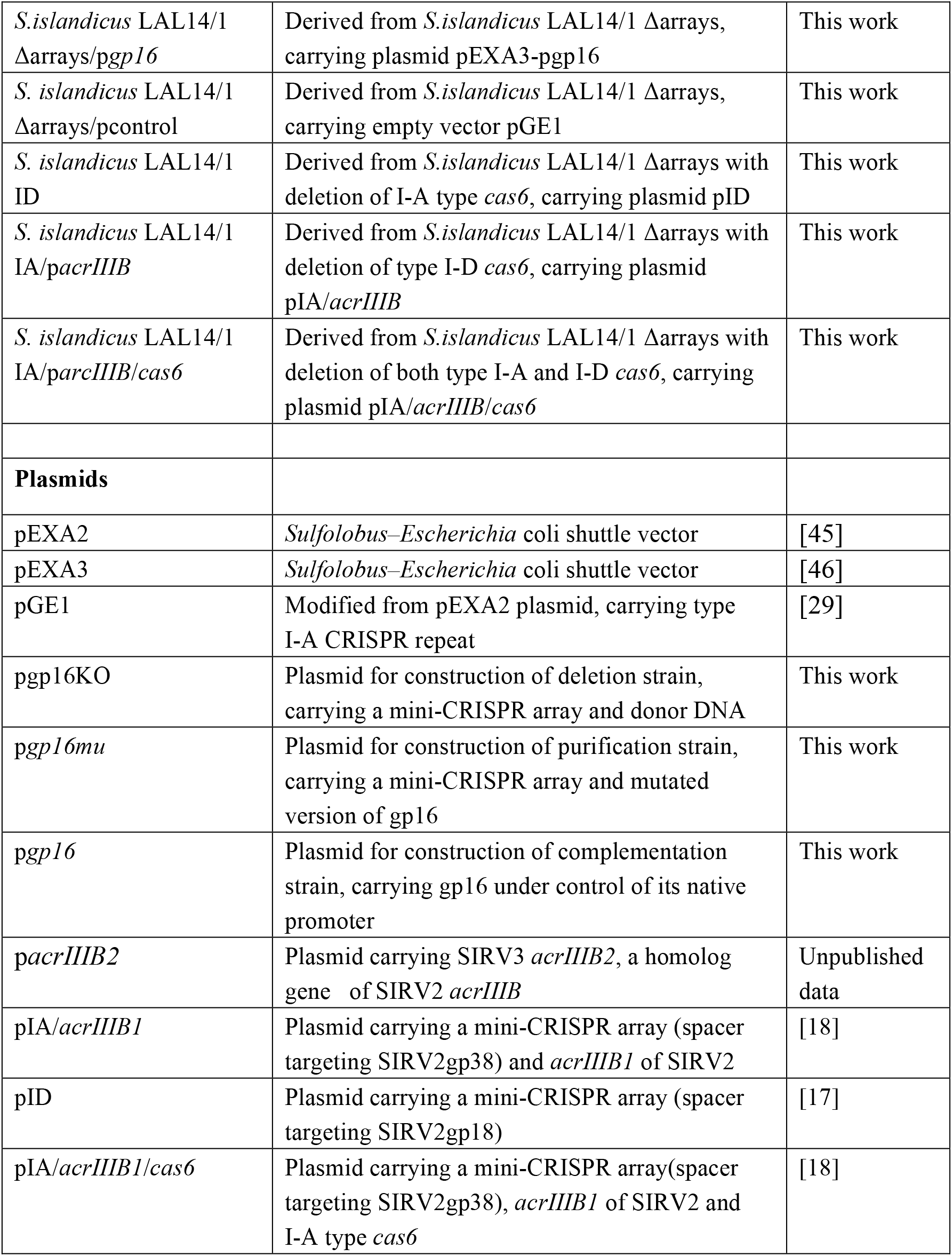
Strains and plasmids.

| Strains | Genotype and features | Source |
| --- | --- | --- |
| *S. islandicus* LAL14/1 | Wild-type strain, ΔpyrEF, with all CRISPR-Cas systems | [44] |
| *S. islandicus* LAL14/1 Δarrays | Derived from *S. islandicus* LAL14/1 without CRISPR arrays | [29] |
| *S. islandicus* LAL14/1 *pacrIIIB2* | Derived from *S. islandicus* LAL14/1, carrying plasmid *pacrIIIB2* | Unpublished data |
| *S. islandicus* LAL14/1 Δarrays/16KO | Derived from *S. islandicus* LAL14/1 Δarrays, carrying plasmid pGE1-16KO | This work |
| *S. islandicus* LAL14/1 Δarrays/pgp16mu | Derived from *S. islandicus* LAL14/1 Δarrays, carrying plasmid pGE1-pgp16mu | This work |
| *S. islandicus* LAL14/1 Δarrays/*pgp16* | Derived from *S. islandicus* LAL14/1 Δarrays, carrying plasmid pEXA3-*pgp16* | This work |
| *S. islandicus* LAL14/1 Δarrays/pcontrol | Derived from *S. islandicus* LAL14/1 Δarrays, carrying empty vector pGE1 | This work |
| *S. islandicus* LAL14/1 ID | Derived from *S. islandicus* LAL14/1 Δarrays with deletion of I-A type *cas6*, carrying plasmid pID | This work |
| *S. islandicus* LAL14/1 IA/*pacrIIIB* | Derived from *S. islandicus* LAL14/1 Δarrays with deletion of type I-D *cas6*, carrying plasmid pIA/*acrIIIB* | This work |
| *S. islandicus* LAL14/1 IA/*parcIIIB/cas6* | Derived from *S. islandicus* LAL14/1 Δarrays with deletion of both type I-A and I-D *cas6*, carrying plasmid pIA/*acrIIIB/cas6* | This work |
| **Plasmids** |  |  |
| pEXA2 | *Sulfolobus*–*Escherichia coli* shuttle vector | [45] |
| pEXA3 | *Sulfolobus*–*Escherichia coli* shuttle vector | [46] |
| pGE1 | Modified from pEXA2 plasmid, carrying type I-A CRISPR repeat | [29] |
| *pgp16*KO | Plasmid for construction of deletion strain, carrying a mini-CRISPR array and donor DNA | This work |
| *pgp16mu* | Plasmid for construction of purification strain, carrying a mini-CRISPR array and mutated version of *gp16* | This work |
| *pgp16* | Plasmid for construction of complementation strain, carrying *gp16* under control of its native promoter | This work |
| *pacrIIIB2* | Plasmid carrying SIRV3 *acrIIIB2*, a homolog gene of SIRV2 *acrIIIB* | Unpublished data |
| pIA/*acrIIIB1* | Plasmid carrying a mini-CRISPR array (spacer targeting SIRV2gp38) and *acrIIIB1* of SIRV2 | [18] |
| pID | Plasmid carrying a mini-CRISPR array (spacer targeting SIRV2gp18) | [17] |
| pIA/*acrIIIB1/cas6* | Plasmid carrying a mini-CRISPR array(spacer targeting SIRV2gp38), *acrIIIB1* of SIRV2 and I-A type *cas6* | [18] |

**Table S4.** Average threshold cycle (Ct) values of qPCR.

| Log<sub>10</sub> dilutions | gp01 | gp26 | gp38 |
| --- | --- | --- | --- |
| -2 | 9.97 | 8.33 | 8.48 |
| -3 | 12.41 | 11.03 | 10.71 |
| -4 | 15.75 | 14.63 | 14.43 |
| -5 | 19.24 | 17.86 | 17.57 |
| -6 | 22.36 | 21.24 | 20.95 |
| -7 | 25.63 | 24.89 | 24.37 |

Virus infected samples
|  |  | gp01 |  |  |  | gp26 |  |  |  | gp38 |  |  |  |
| --- | --- | --- | --- | --- | --- | --- | --- | --- | --- | --- | --- | --- | --- |
| min p.i. | Diluted fold | pgp16+Δgp16-a |  | pcontrol+Δgp16-a |  | pgp16+Δgp16-a |  | pcontrol+Δgp16-a |  | pgp16+Δgp16-a |  | pcontrol+Δgp16-a |  |
| 0 | 10 | 19.93 | 19.92 | 19.40 | 19.56 | 20.02 | 19.96 | 19.49 | 19.81 | 19.45 | 19.68 | 18.95 | 19.32 |
| 30 | 100 | 21.45 | 21.82 | 21.33 | 21.34 | 21.14 | 21.47 | 21.12 | 21.27 | 20.84 | 21.35 | 20.75 | 20.78 |
| 60 | 100 | 20.36 | 20.41 | 21.69 | 22.09 | 20.96 | 21.08 | 21.26 | 21.48 | 20.75 | 20.77 | 20.94 | 21.39 |
| 120 | 100 | 18.78 | 18.85 | 21.19 | 21.35 | 19.44 | 19.56 | 20.79 | 20.52 | 19.30 | 19.50 | 20.42 | 20.49 |
| 240 | 1000 | 20.40 | 19.88 | 21.09 | 20.49 | 20.09 | 19.29 | 20.67 | 19.83 | 19.72 | 19.48 | 20.41 | 19.99 |
| 360 | 1000 | 19.83 | 19.48 | 21.59 | 20.96 | 19.88 | 19.34 | 21.41 | 20.51 | 20.03 | 19.58 | 21.01 | 20.34 |
| 480 | 1000 | 19.46 | 19.39 | 20.18 | 20.39 | 18.95 | 19.08 | 20.00 | 20.10 | 18.86 | 18.99 | 19.62 | 20.05 |

**Table S5.** PCR programs.

| Primer pair | gp26 |  |  | gp16 mutant virus |  |  | gp16 wild-type virus |  |  |
| --- | --- | --- | --- | --- | --- | --- | --- | --- | --- |
| Stage | Temperature (°C) | Duration | Number of cycles | Temperature (°C) | Duration | Number of cycles | Temperature (°C) | Duration | Number of cycles |
| Initial Denaturation | 95°C | 180 s | × 1 | 95°C | 180 s | × 1 | 95°C | 180 s | × 1 |
| Denaturation | 95°C | 15 s | × 27 | 95°C | 15 s | × 30 | 95°C | 15 s | × 30 |
| Annealing | 57°C | 15 s |  | 54.3°C | 15 s |  | 50.9°C | 15 s |  |
| Extension | 72°C | 30 s |  | 72°C | 30 s |  | 72°C | 30 s |  |
| Final Extension | 72°C | 300 s | × 1 | 72°C | 300 s | × 1 | 72°C | 300 s | × 1 |

